## Supplementary Material for "Partitioning gene-based variance of complex traits by gene score regression"

#### S1 Figures

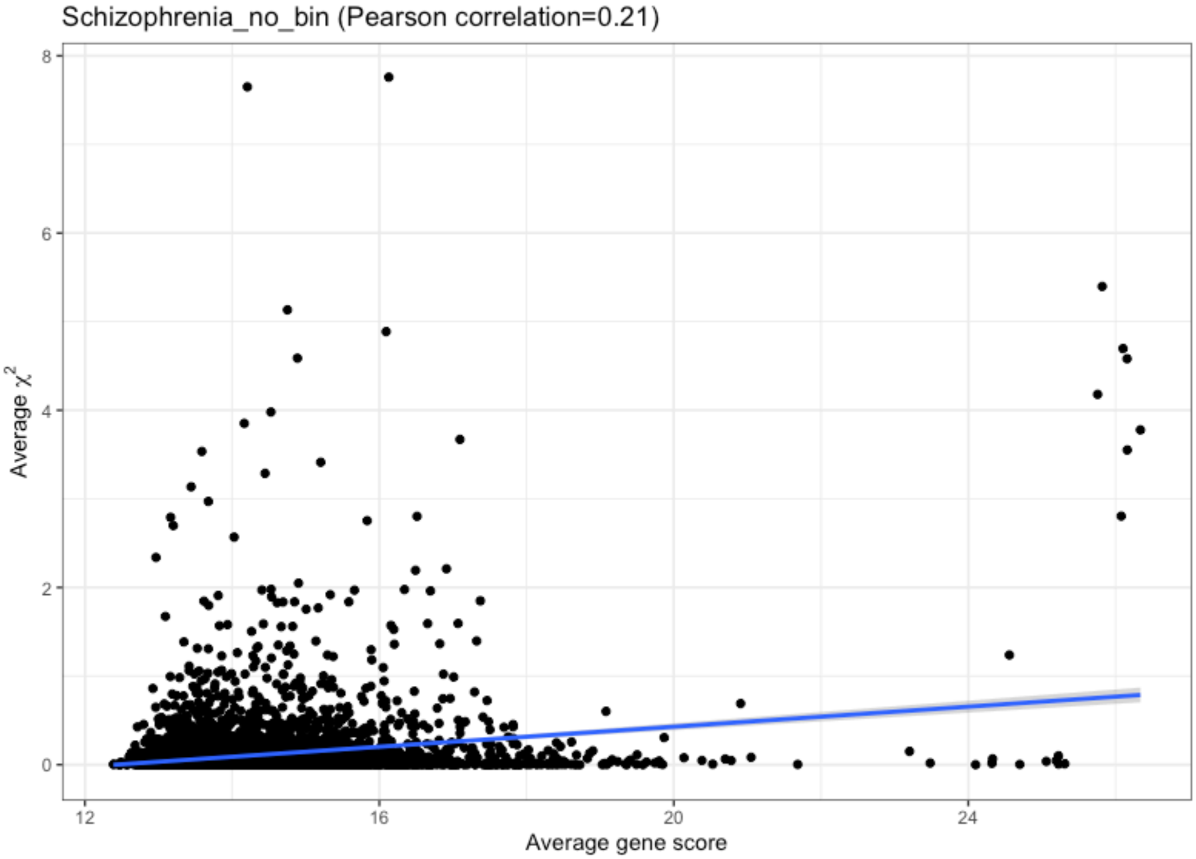


Figure 1: Gene scores were correlated with marginal TWAS chi-square statistics of Schizophrenia. An binned version is supplied in Figure 2. The fitted slope is 0.044 (SD = 0.0039) compared to 0.056 (SD = 0.0035), fitted by binned gene scores and $\chi^2$.

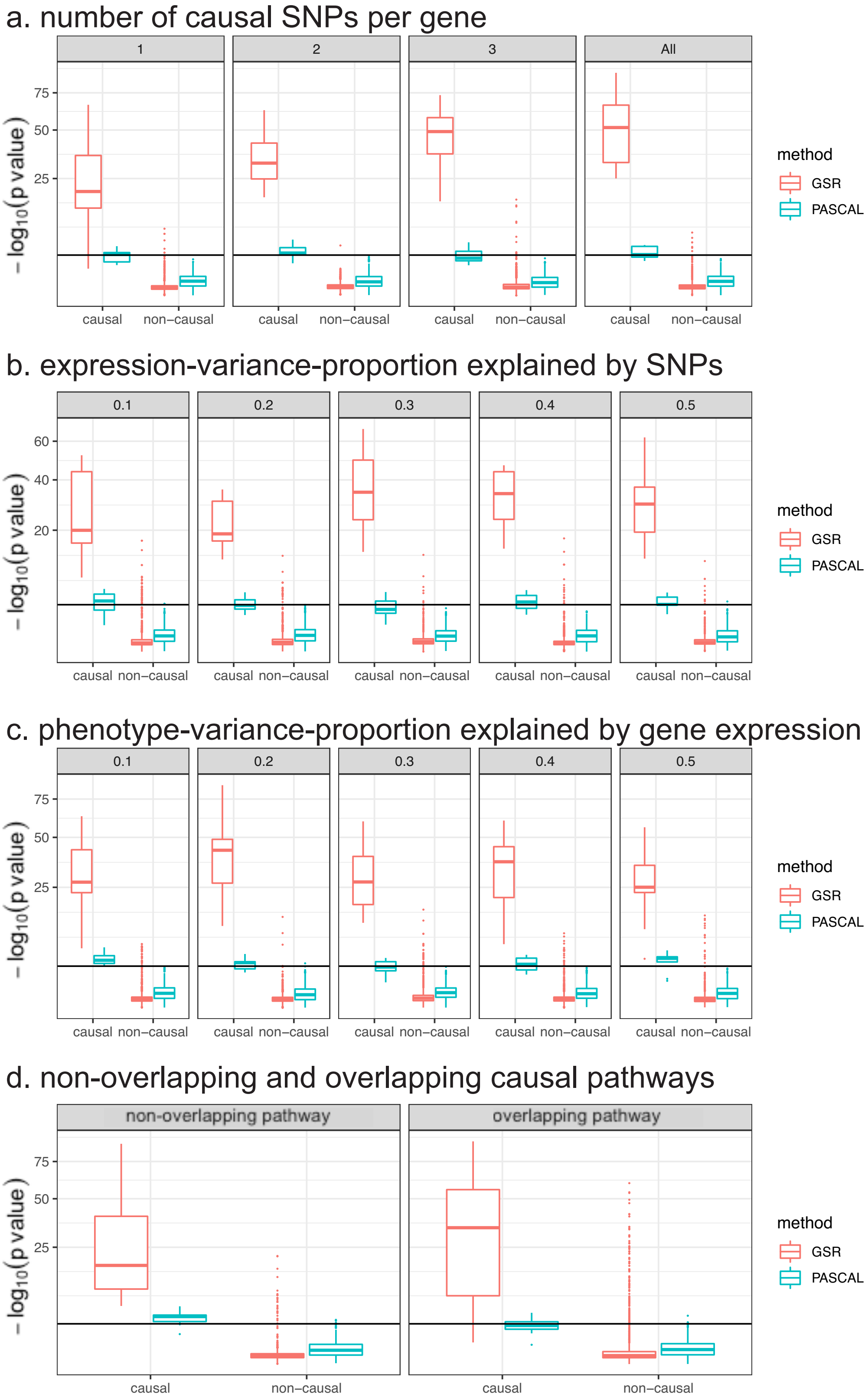


Figure 2: Simulation to evaluate methods in various simulation settings. We compared GSR and PASCAL [1] working with (a) different number of causal SNP per gene (b) different SNP-gene heritability; (3) different gene-phenotype variance explained; and (d) different overlapping causal pathway. For visibility of the large p-values, we plot the y-axis on the scale of the squared root of $-log_{10}$(p-values).

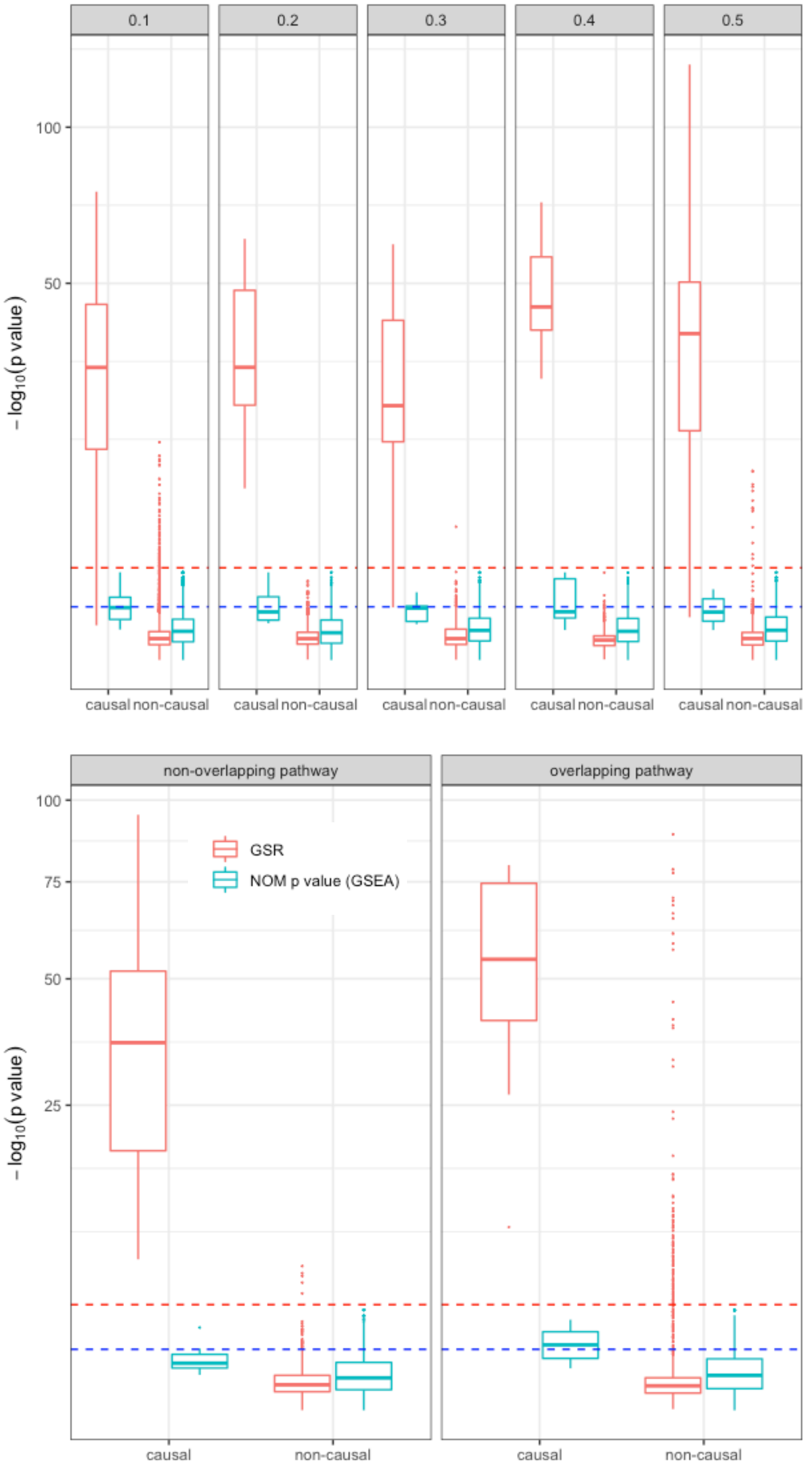


Figure 3: Comparison of pathway enrichment between GSR and GSEA [2]. Pathway enrichment was inferred using observed gene expression.

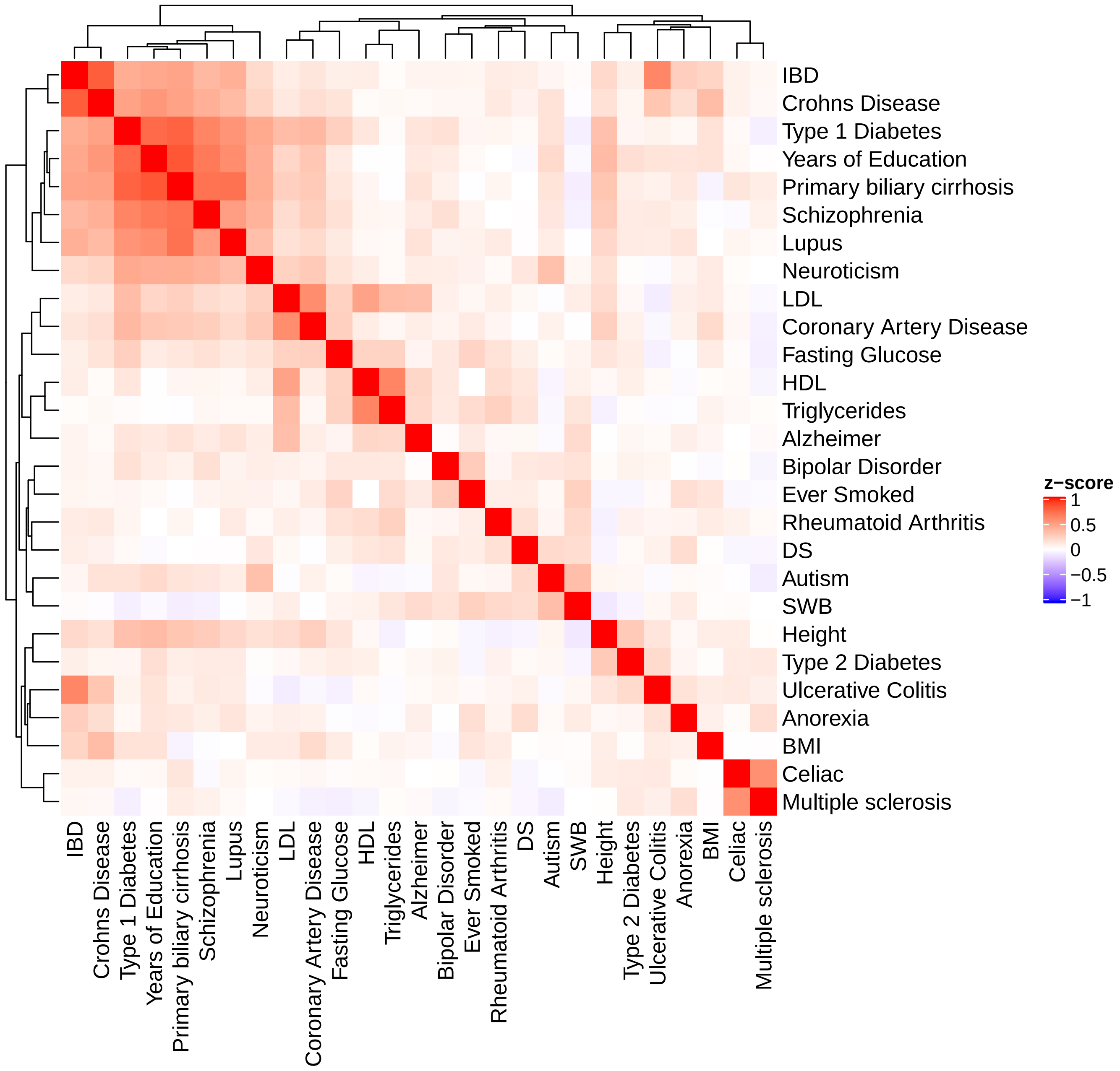


Figure 4: Gene set enrichments suggest shared genetic and biological pathways by related complex traits

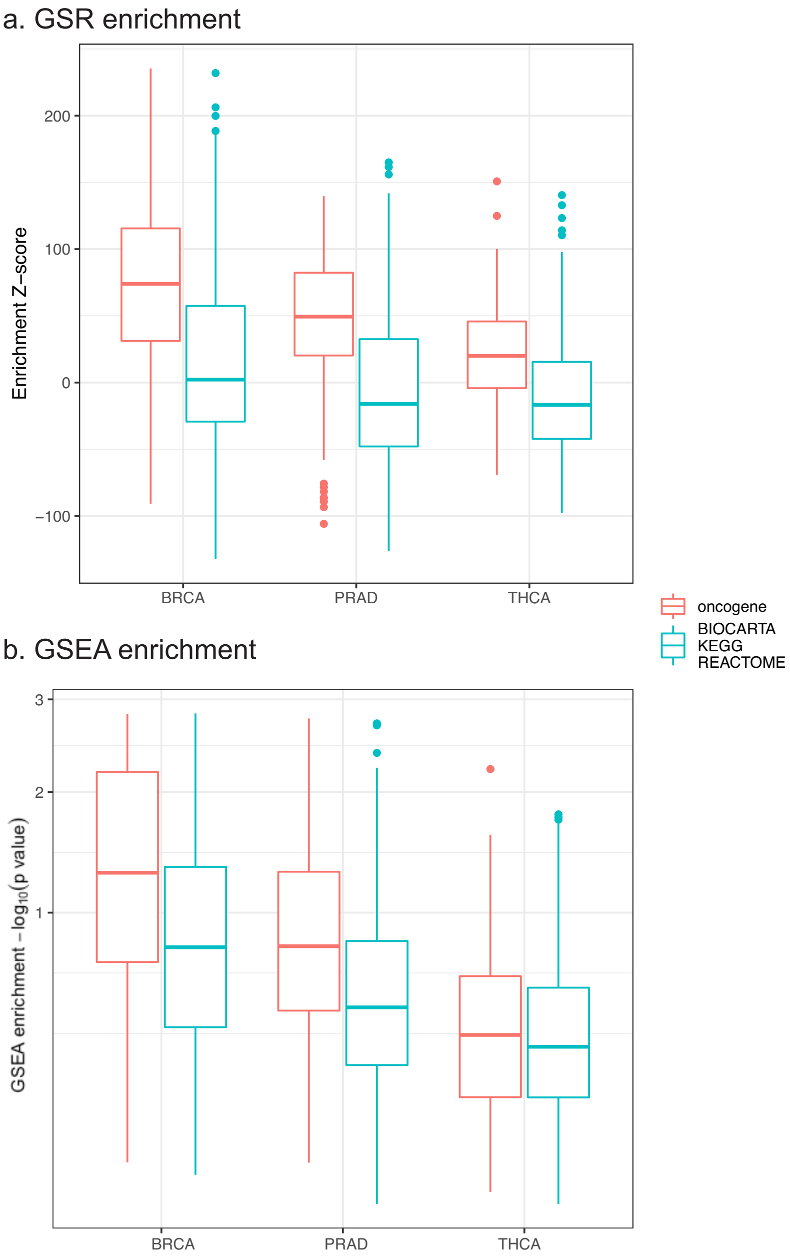


Figure 5: Tumour-specific gene set enrichment. GSR and GSEA were performed separately on the processed gene expression data of BRCA, PRAD and THCA samples versus normal samples from TCGA and GTEx. We tested the enrichments for oncogenic signatures as well as 1,050 gene sets obtained from BIOCARTA, KEGG and REACTOME. We then separately plotted the enrichment Z-scores for oncogenic gene sets and the general gene sets across the three tumour types.

### S2 Supplementary Tables

Table 1: Gene set enrichments on select traits. The enrichments were applied to three gene sets: (1) 4,360 GO Biology Processes terms from MSigDb; (2) combined 1,051 BIOCARTA, KEGG, REACTOME gene sets from MSigDb; and (3) 207 gene sets each derived from GTEx and Franke tissue/cell-specifically expressed genes.

| trait | gene set | p-value | t-value |
|---|---|---|---|
| HDL | REACTOME_CHYLOMICRON_MEDIATED_LIPID_TRANSPORT | 3.23e-151 | 26.98 |
| HDL | GO_NEUTRAL_LIPID_CATABOLIC_PROCESS | 1.70e-132 | 25.14 |
| HDL | GO_TRIGLYCERIDE_CATABOLIC_PROCESS | 1.70e-132 | 25.14 |
| HDL | GO_MACROMOLECULAR_COMPLEX_REMODELING | 5.82e-131 | 24.98 |
| HDL | GO_ACYLGLYCEROL_HOMEOSTASIS | 1.11e-113 | 23.17 |
| HDL | REACTOME_LIPOPROTEIN_METABOLISM | 3.99e-112 | 23.00 |
| HDL | REACTOME_LIPID_DIGESTION_MOBILIZATION_AND_TRANSPOR | 5.93e-96 | 21.17 |
| HDL | GO_PROTEIN_LIPID_COMPLEX_SUBUNIT_ORGANIZATION | 2.58e-90 | 20.51 |
| HDL | REACTOME_PACKAGING_OF_TELOMERE_ENDS | 2.65e-42 | 13.74 |
| HDL | KEGG_PPAR_SIGNALING_PATHWAY | 1.72e-32 | 11.94 |
| Lupus | GO_POSITIVE_REGULATION_OF_INTERFERON_ALPHA_PRODUCT | 7.06e-150 | 26.85 |
| Lupus | GO_POSITIVE_REGULATION_OF_INTERFERON_BETA_PRODUCTI | 1.46e-101 | 21.82 |
| Lupus | GO_RESPONSE_TO_MURAMYL_DIPEPTIDE1 | 2.26e-96 | 21.22 |
| Lupus | GO_POSITIVE_REGULATION_OF_INTERLEUKIN_12_PRODUCTIO | 1.39e-78 | 19.05 |
| Lupus | GO_REGULATION_OF_INTERFERON_ALPHA_PRODUCTION | 1.38e-72 | 18.27 |
| Lupus | REACTOME_INTERFERON_GAMMA_SIGNALING | 2.90e-65 | 17.28 |
| Lupus | REACTOME_INTERFERON_ALPHA_BETA_SIGNALING | 1.30e-29 | 11.36 |
| Lupus | KEGG_TOLL_LIKE_RECEPTOR_SIGNALING_PATHWAY | 1.70e-24 | 10.26 |
| Lupus | REACTOME_INTERFERON_SIGNALING | 1.18e-13 | 7.44 |
| Lupus | REACTOME_CYTOKINE_SIGNALING_IN_IMMUNE_SYSTEM | 5.11e-07 | 5.03 |
